# Defining the domain organization of K1-like killer toxins

**DOI:** 10.1101/2024.10.11.617966

**Authors:** Sarah A. Coss, Lynda E. Clayton, F. Marty Ytreberg, Paul A. Rowley

## Abstract

Antifungal killer toxins produced by yeasts undergo post-translational processing to enable the formation of mature and active toxins. The preprocessed, immature K1 killer toxin from *Saccharomyces cerevisiae* has four different domains (delta, alpha, gamma, and beta) that are cleaved to produce a mature toxin consisting of a disulfide-linked alpha and beta heterodimer. K1 homologs from various yeast species of the Saccharomycotina have been previously identified by primary sequence homology. Primary sequence analysis has identified putative cleavage sites across all homologs, with some conservation with cleavage sites identified in K1 and the likely importance of the conserved carboxypeptidases Kex1 and Kex2. The identification of sites of proteolytic processing has enabled an accurate definition of the domain boundaries of delta, alpha, gamma, and beta domains in all K1 homologs. Functional assays revealed that many K1 homologs contain cytotoxic alpha domains. Interestingly, several killer toxins that lacked measurable antifungal activity still exhibited toxic alpha domains when expressed independently. These findings suggest that alpha domain toxicity is conserved across K1 homologs, and that variations in post-translational processing and domain interactions may influence full-length toxin functionality. This work provides insights into the evolutionary conservation and diversity of yeast killer toxins.

## Introduction

Many yeast species produce extracellular proteinaceous “killer” toxins thought to aid niche competition. Killer toxins are secreted protein toxins that are post-translationally processed by disulfide bond formation and proteolysis to form a mature extracellular toxin. Of all the killer toxins produced by yeasts, the K1 toxin produced by the brewer’s yeast *Saccharomyces cerevisiae* is the most well-studied. The immature pre-processed (prepro) K1 is organized from the N-terminus into four domains named delta, alpha, gamma, and beta (Figure 1). The first step in the post-translational processing of K1 is export to the endoplasmic reticulum (ER) and cleavage of the N-terminal signal peptide after amino acid 26, resulting in processed (pro) K1. In the lumen of the endoplasmic reticulum, disulfide bonds form between cysteine residues and are predicted to link the alpha and beta domains. Subsequent export of pro-K1 to the Golgi leads to cleavage after R44, R149, and R233 by the proprotein convertase and endoprotease Kex2 to remove the delta and gamma domains. The location of Kex2 cleavage occurs after a dibasic motif with a basic residue (arginine or lysine) at the P2 position, followed by arginine at the P1 position. Residues after the cleavage site have also been shown to influence the efficiency of cleavage, with negatively charged and small amino acids preferred at sites P1’, P2 ‘, and P4’. Moreover, a dibasic motif alone is not sufficient to establish cleavage because extended sequence context and structural accessibility influence the extent of Kex2 processing. The carboxypeptidase Kex1 processes the Kex2-cleaved alpha domain by the removal of two arginine residues at the C-terminal end of the K1 alpha domain. After processing, the resulting mature K1 toxin that is exported from the cell is a heterodimer with an alpha and beta subunit linked by at least one intramolecular disulfide bond.

**Figure 1.**
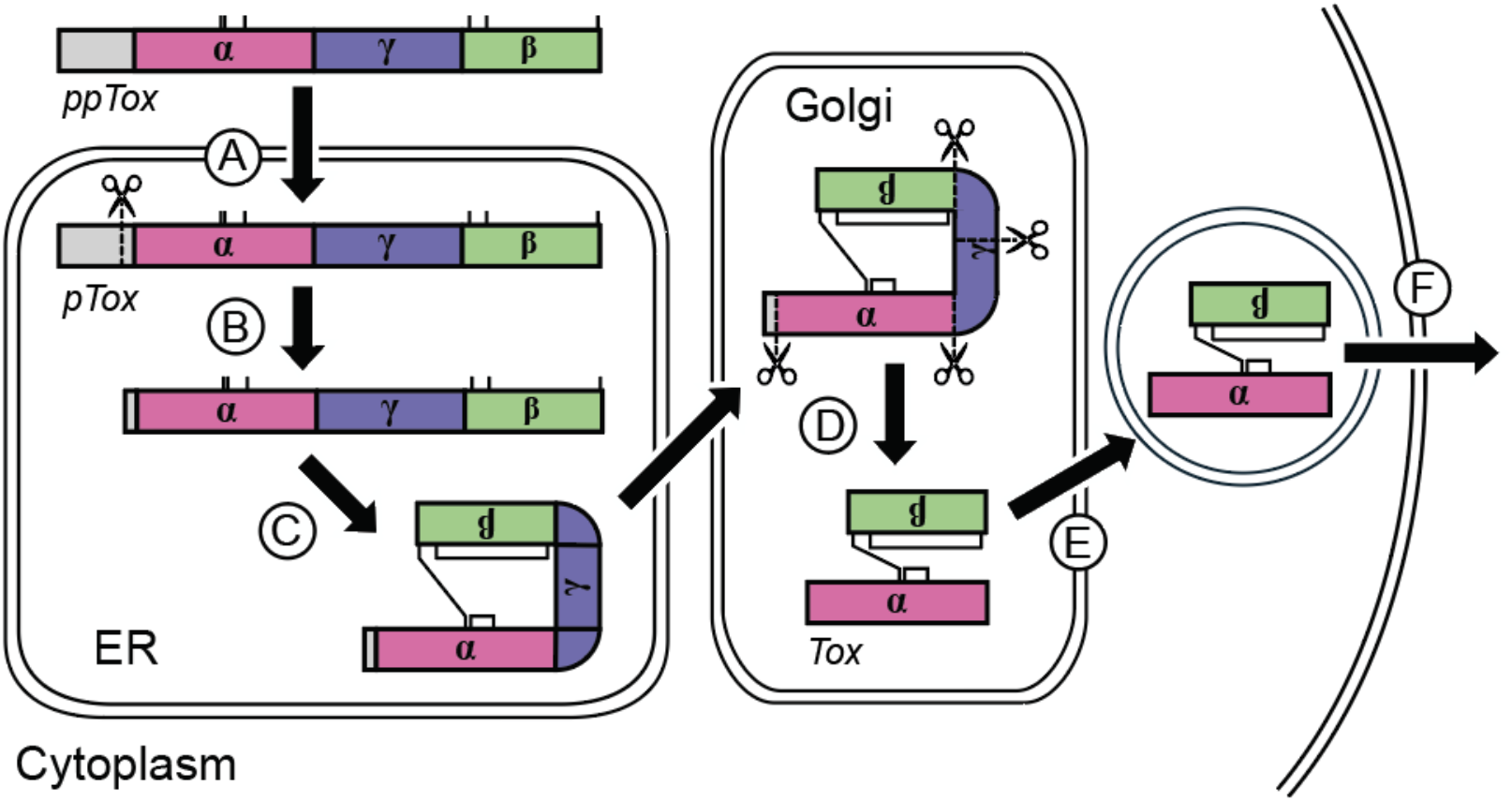
A model of K1 killer toxin processing. Immature prepro-K1 (ppTox) encodes a signal sequence for endoplasmic reticulum (ER) export and four domains: delta, alpha, gamma, and beta. (A) After translocation to the ER, (B) the signal sequence is cleaved to form pro-K1 (pTox), after which (C) disulfide bonds are formed within and between the alpha and beta domains. After transport to the Golgi network, (D) the carboxypeptidas Kex1 and endopeptidase Kex2 cleave pTox, releasing the delta and gamma domains. (E) Mature K1 toxin is packaged into export vesicles and (F) exported from the cell.

The mechanism of K1 attack targets the plasma membrane, where it disrupts ion homeostasis by triggering potassium efflux. The mechanism of membrane attack by K1 remains to be fully determined, but is likely based on channel formation. The secondary structure of the K1 alpha domain is predominantly alpha-helical with hydrophobic regions and the potential to form transmembrane helices, suggesting a direct interaction with membranes. Expression of the K1 alpha domain (K1-α) by *S. cerevisiae* results in cell death in a receptor-independent manner. A family of K1 homologs has been found in other yeast species of the Saccharomycotina subphylum with similar length to prepro-K1 (∼340 amino acids). Amino acid sequence identity between the alpha domains of these homologs and K1 ranges between 16-28%, and is the most conserved domain of these proteins. Ectopic expression of K1 homologs in *S. cerevisiae* has shown that these genes encode active killer toxins (K1L (*Saccharomyces paradoxus*), *KKT1* (*Pichia membranifaciens*), *KKT2* (*Kazachstania africana*), and *KKT1* (*Nauvomazyma dairenensis*). Although a subset of these K1-like proteins have been confirmed as active toxins, it is unclear whether they share common post-translational processing and intoxication mechanisms with K1.

This manuscript describes a functional analysis of K1 homologs to define domain boundaries by examining post-translational modification via proteolytic cleavage. Using molecular modeling and empirical assays, we predicted the domain boundaries of delta, alpha, gamma, and beta for 13 K1 homologs. Cloning and expression of alpha domains confirmed that, similar to K1, alpha domains from diverged toxins were lethal to *S. cerevisiae*, including domains cloned from toxins that lacked prior antifungal activity, suggesting that these toxins are potentially active. The dependency of posttranslational processing on the Kex proteases was also confirmed, and the importance of the specific dibasic sites was validated by mutagenesis. This was further validated by detecting partially processed intermediates after the mutation of dibasic Kex cleavage sites. These data show that despite divergence in protein primary sequence, the domain order and organization, as well as tertiary structures, for most K1-homologs are conserved.

## Results

### K1 homologs require KEX1 and KEX2 for their antifungal activity

The proteases Kex1 and Kex2 are required for posttranslational modification and maturation of K1, but it is unknown if K1 homologs require the same enzymes. Killer toxins from *Pichia membranifaciens* (PmKKT1), *Kazachstania africana* (KaKKT2), and *Naumovozyma dairenensis* (NdKKT1) under the control of a galactose-inducible promoter were expressed in wild-type *S. cerevisiae* and strains lacking the genes *KEX1* or *KEX2*. Killer toxin-expressing strains were plated on killer assay media containing dextrose or galactose as the sole carbon source to control killer toxin expression. As expected, no killer toxins were produced when genes were not expressed on dextrose-containing media (Figure 2A). When strains were grown with galactose, zones of growth inhibition were observed around wild-type *S. cerevisiae* expressing K1, PmKKT1, KaKKT2, and NdKKT1, indicating the inducible expression of these antifungal killer toxins. Overexpression of K1 and KaKKT2 did not result in severe growth defects, whereas PmKKT1 and NdKKT1 caused a modest drop in growth rate on agar (Figure 2B). The deletion of *KEX1* (*kex1Δ*) or *KEX2* (*kex2Δ*) resulted in a loss of antifungal activity for K1 and its homologs (Figure 2A). While the *kex2Δ* deletion resulted in the complete loss of killer toxin production for all toxins, a small zone of growth inhibition was observed in the *kex1Δ* mutant expressing KaKKT2. These data show that *KEX1* and *KEX2* are essential for producing fully active K1-like killer toxins, as has been shown for K1.

**Figure 2.**
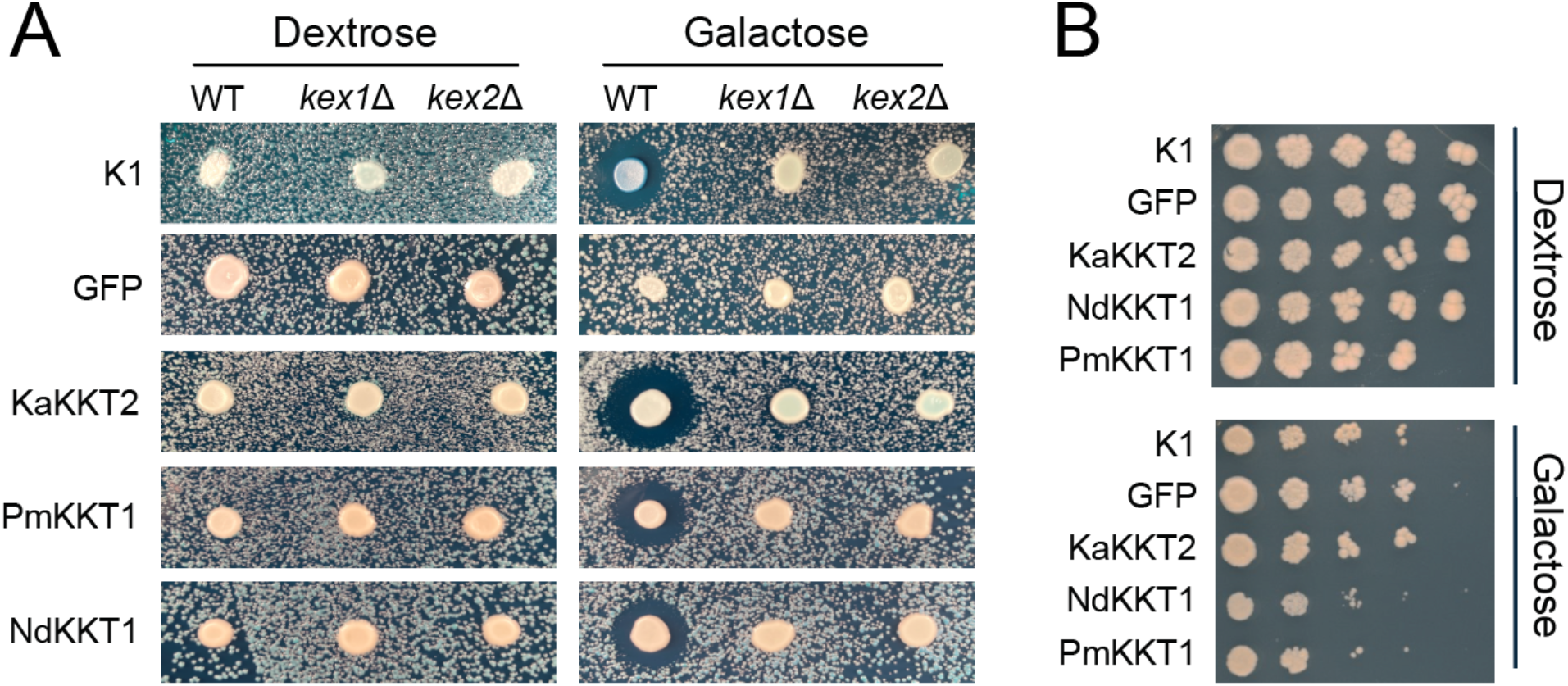
Killer toxin activity is dependent on the presence of *KEX1* and *KEX2*. Plasmid-encoded K1 homologs or GFP were cloned under a galactose-inducible promoter in the non-killer laboratory yeast *S. cerevisiae* (BY4741) with or without deletions of *KEX1* (*kex1Δ*) or *KEX2* (*kex2Δ*). (A) Killer toxin expression was induced by galactose or suppressed by dextrose on methylene blue agar at pH 4.6 to measure killer toxin activity against a lawn of *K. africana*. (B) Killer toxin expressing yeast were also plated on YPD or YPG to observe any growth defect as a result of toxin overexpression. All images are representative of at least n>3 replicates.

### Dibasic sequences are conserved in K1 homologs and define domain boundaries

As *KEX1* and *KEX*2 are required for the antifungal activities of the K1 homologs PmKKT1, KaKKT2, and NdKKT1, we speculated that these enzymes are required for the post-translational modification of these toxins. K1 cleavage by Kex proteases occurs at four locations, separating the alpha and beta domains (which constitute the mature toxin) from the gamma and delta domains (Figure 3A). An additional cleavage motif in the central region of the K1 gamma domain also reduces K1 toxicity but not immunity when mutated. To determine whether K1 homologs contain potential Kex cleavage sites, we performed a multiple sequence alignment of 13 K1 homologs to identify conserved basic amino acids aligned with known K1 cleavage sites. Moreover, secondary structure predictions were also included for each toxin to define the structural context of these cleavage sites.

**Figure 3.**
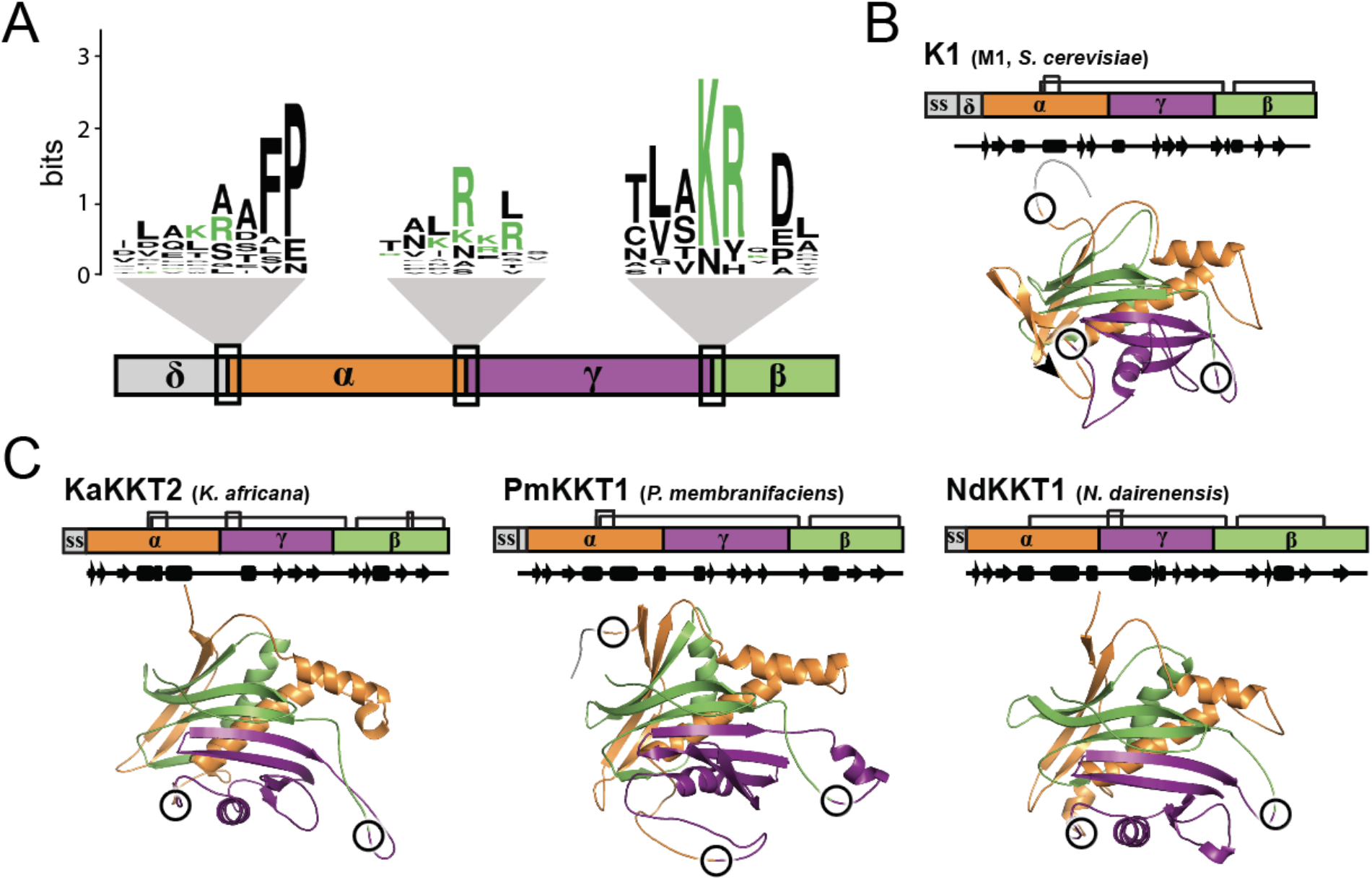
Predicted domain boundaries in K1 homologs based on primary, secondary, and tertiary structure conservation. (A) Logo of the primary sequence surrounding predicted Kex cleavage sites in K1 and its homologs. (B) Tertiary structure prediction and known domain boundaries of the K1 killer toxin encoded on the M1 satellite of *S. cerevisiae*. (C) Tertiary structure prediction and predicted domain boundaries of the killer toxins KaKKT2, PmKKT1, and NdKKT1. Linear domain diagrams show relative domain length and position, as well as disulfide bonds. The tertiary structure is colored to match the linear domain representation. Circles on tertiary structures represent the location of known and putative Kex protease cleavage sites.

Signal sequence cleavage was predicted using SignalP and TargetP (Table 1). Of the sites predicted there was agreement between the two methods for 71% of the cleavage sites (Table S1), with only TargetP correctly predicting the known signal sequence cleavage point of K1. TpKKT4 was the only putative toxin that did not have a predicted signal sequence by both methods (Figure 2). Other sites of posttranslational cleavage by Kex proteases were inferred by sequence conservation with K1. This approach was complicated by a lack of alignment due to the low sequence homology across homologs and multiple basic residues with potential cleavage sites. Therefore, predictions were made on the positioning of the basic residues relative to the predicted secondary structure of K1-homologs (Figure 3C and S1). The first cleavage site in K1 is at the boundary of the delta and alpha domains (Kex-δ/α) after the basic residue R44 (PR↓E45) (Figure 3B and Table 1) This residue is positioned after the predicted signal sequence but before the first secondary structure element of the alpha domain. Of the 13 K1 homologs, four have the preferred KR dibasic motif for carboxypeptidase cleavage at Kex-δ/α, NdKKT1 has a monobasic motif (IR↓L26), and NcKKT1 had K39 aligned close to the K1 cleavage site, but no arginine that would direct Kex2 cleavage (Figure 3). Of the remaining nine homologs that lack obvious Kex cleavage sites at the δ/α boundary, seven had a predicted signal sequence cleavage site at a location similar to K1 Kex-δ/α (Table 1). This suggested that, in these homologs, signal-peptide cleavage separates the delta and alpha domains. Only TpKKT4 lacked both Kex cleavage sites and a recognizable signal sequence. The two mechanisms of removing the delta domain meant that the consensus of the putative cleavage site was not dominated by the typical dibasic motif (Figure 2A).

**Table 1.**
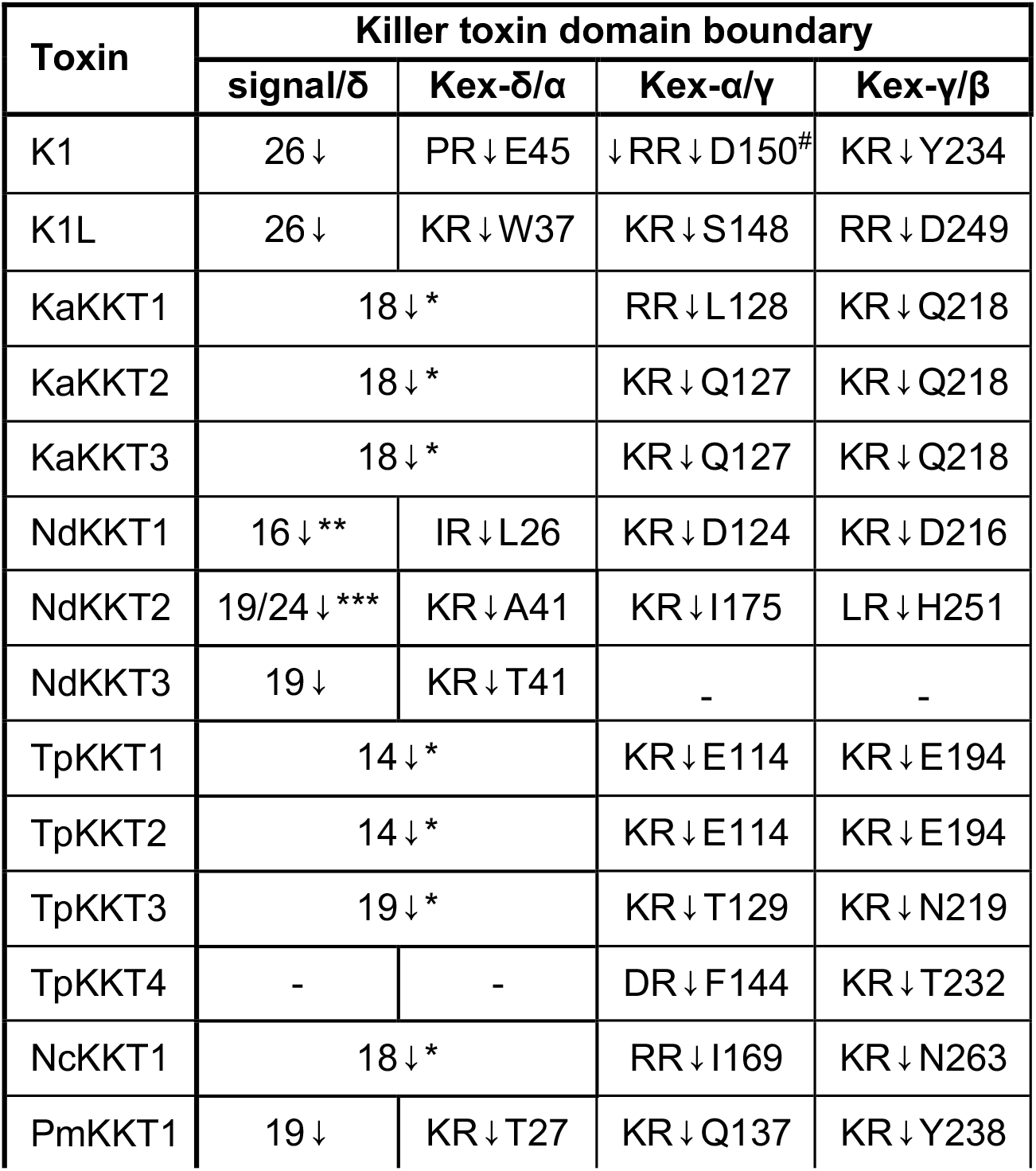
Location of predicted domain boundaries of K1-like killer toxins. The dibasic motif prior to each predicted Kex cleavage point is shown, with the cleavage point represented as an arrow (↓) followed by the number of the amino acid on the N-terminus of the domain boundary. - no cleavage site identified. Signal/δ indicates the cleavage point of the signal peptidase. *Domain boundary at δ/α cleaved by signal peptidase. **Signal peptidase cleavage site predicted only by TargetP. ***Two different signal peptidase cleavage sites predicted by SignalP and TagetP. ^#^Kex2 cleavage has been shown experimentally to remove the dibasic motif at this cleavage point.

| Toxin | Killer toxin domain boundary |  |  |  |
| --- | --- | --- | --- | --- |
| | signal/$\delta$ | Kex-$\delta/\alpha$ | Kex-$\alpha/\gamma$ | Kex-$\gamma/\beta$ |
| K1 | 26↓ | PR↓E45 | ↓RR↓D150<sup>#</sup> | KR↓Y234 |
| K1L | 26↓ | KR↓W37 | KR↓S148 | RR↓D249 |
| KaKKT1 | 18↓* |  | RR↓L128 | KR↓Q218 |
| KaKKT2 | 18↓* |  | KR↓Q127 | KR↓Q218 |
| KaKKT3 | 18↓* |  | KR↓Q127 | KR↓Q218 |
| NdKKT1 | 16↓** | IR↓L26 | KR↓D124 | KR↓D216 |
| NdKKT2 | 19/24↓*** | KR↓A41 | KR↓I175 | LR↓H251 |
| NdKKT3 | 19↓ | KR↓T41 | - | - |
| TpKKT1 | 14↓* |  | KR↓E114 | KR↓E194 |
| TpKKT2 | 14↓* |  | KR↓E114 | KR↓E194 |
| TpKKT3 | 19↓* |  | KR↓T129 | KR↓N219 |
| TpKKT4 | - | - | DR↓F144 | KR↓T232 |
| NcKKT1 | 18↓* |  | RR↓I169 | KR↓N263 |
| PmKKT1 | 19↓ | KR↓T27 | KR↓Q137 | KR↓Y238 |

**Figure S1.**
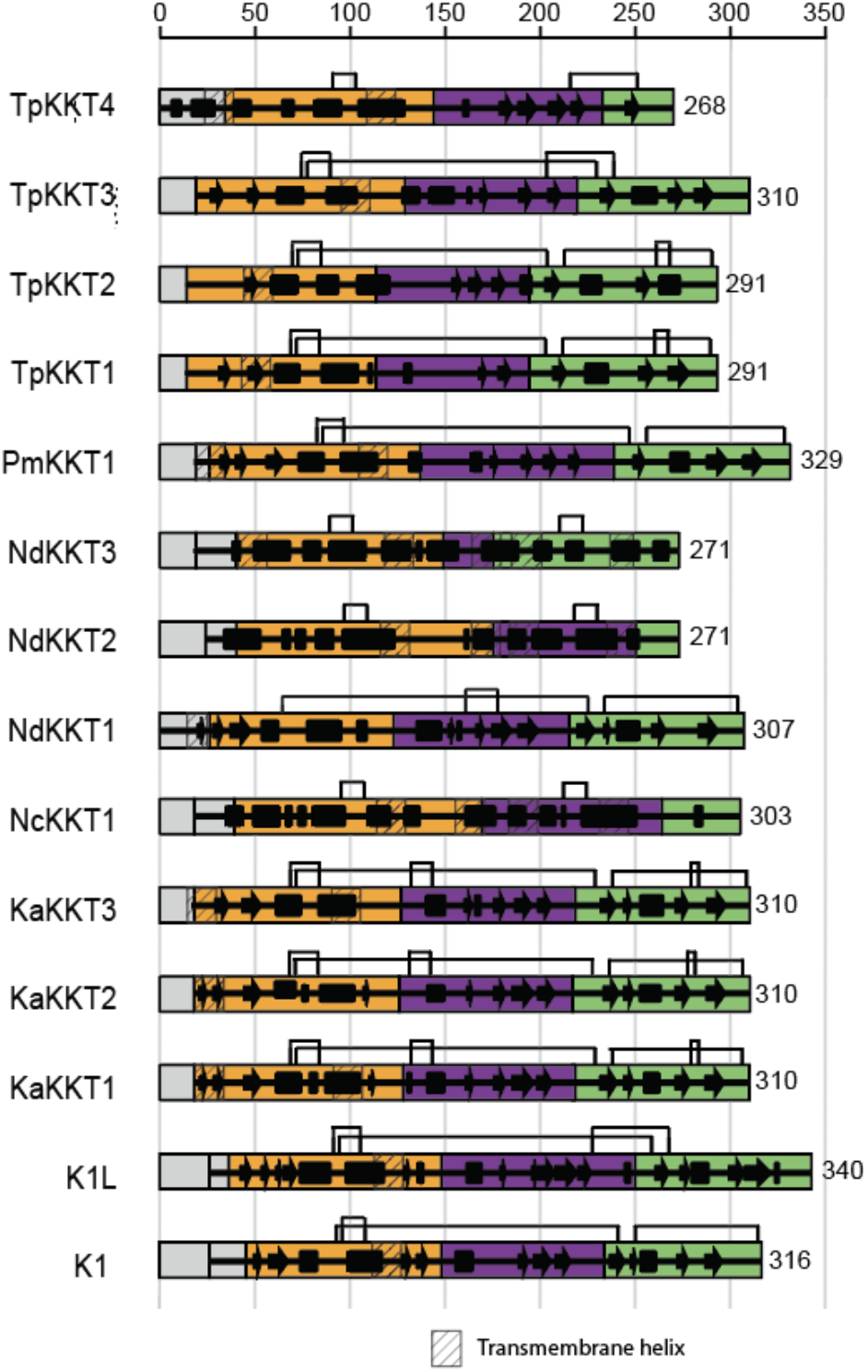
Secondary structure predictions and domain boundaries in K1 and K1 homologs. Secondary structure prediction and predicted domain boundaries of K1 and its homologs. Linear domain diagrams show relative domain length and position, and disulfide bonds. The tertiary structure is colored (from the left) according to delta (grey), alpha (orange), gamma (purple), and beta (green). Sequences predicted to be transmembrane are shown as hatched boxes.

The second conserved dibasic site at the boundary of the alpha and gamma domains (Kex-α/γ) is present after the beta sheet four and before helix three in K1. As with the first cleavage site, despite overall poor sequence conservation, 9/13 had at least two consecutive basic residues within one residue of the aligned K1 Kex-α/γ. 6/13 of the putative cleavage sites were KR, and 3/13 had a run of 3-4 consecutive basic residues (RKR or RKRR). The consensus sequence for Kex-α/γ was determined to be LRxL/R. While NdKKT3 lacked arginine residues at the α/γ boundary, NdKKT2 and NcKKT1 had dibasic motifs that were aligned away from the canonical K1 Kex-α/γ site. This alternative positioning lengthened the alpha domains of NdKKT2 and NcKKT1 by 29 and 24 residues relative to K1, respectively.

The third conserved dibasic site at the boundary between the gamma and beta domains (Kex-γ/β) is positioned on an unstructured loop in K1. The majority of the predicted Kex-γ/β sites in K1 homologs (9/13) were KR, and one was RKR. The consensus sequence at Kex-γ/β was determined to be “LAKRxDxP”, where the basic residues at P2 (K) and P1 (R) are conserved in 10/13 of K1-like killer toxins, and the P2’ sequence was negatively charged with an aspartate residue. The resulting beta domains had similar patterns of secondary structure with a beta-strand/alpha-helix/beta-strand organization in 9/13 homologs similar to K1, despite little to no sequence homology. The putative toxins that lacked K1-like cleavage sites or had unusually short beta domains lacked this secondary structure pattern (NcKKT1, NdKKT2, NdKKT3, and TpKKT4). The dibasic cleavage that is centrally located in the gamma domain of K1 was only identified in 5/13 of the K1 homologs and was also positioned centrally; as this cleavage site does not represent a recognized domain boundary, it was not analyzed further. Overall, domain boundaries defined by the conservation and positioning of basic residues relative to K1 resulted in similarly sized domains for the majority of K1 homologs. The alpha domain was the largest, with an average length of 113 amino acids (SD +/-14), and is the most highly conserved domain by sequence identity. The gamma and beta domains were smaller, with average lengths of 89 and 79 amino acids, respectively (Table S2).

To validate the accessibility of monobasic or dibasic residues for posttranslational cleavage, tertiary structure molecular modeling was performed using AlphaFold2 to obtain structure predictions for K1-like homologs. AlphaFold2 assessed confidence in the predicted tertiary structures by assigning a predicted Local Distance Difference Test (pLDDT) score to each residue of each K1-like toxin (Figure S2 and Table S3). Scores for this test can range from 0 to 100, indicating the lowest to highest confidence. The highest pLDDT score among all models was 72.4, the lowest was 37.2, and the average across all predicted structures was 56.3. These models had similar pLDDT scores to those previously published for AlphaFold models of K1 (global, 51.35; min, 25.24; max, 72.3) and K1L (global, 62.93; min, 26.21; max, 87.72). The structures with the lowest pLDDT scores, NdKKT2 (38.2), NdKKT3 (37.2), and NcKKT1 (39.3), had some of the lowest sequence identity to K1 (<12%) and the lowest-confidence structural models. Despite the lower pLDDT scores, most models showed structural homology to K1, with an average root mean square deviation (RMSD) of 6 Å. The AlphaFold models also underwent 100 ns molecular dynamics (MD) simulations, with all models showing stable RMSD values after the first ∼25 ns (Figure S3). RMSF plots showed that the killer toxin models were flexible during the simulation, primarily at exposed loops (Figure S4). Residues in Ramachandran-favored regions remained largely unchanged after simulation, with an average of 89.7% of residues positioned in favored regions pre-simulation and 89.9% of residues in favored regions post-simulation, with a standard deviation of 4.1 (Table S4 <u>KKTs tables</u>).

**Figure S2.**
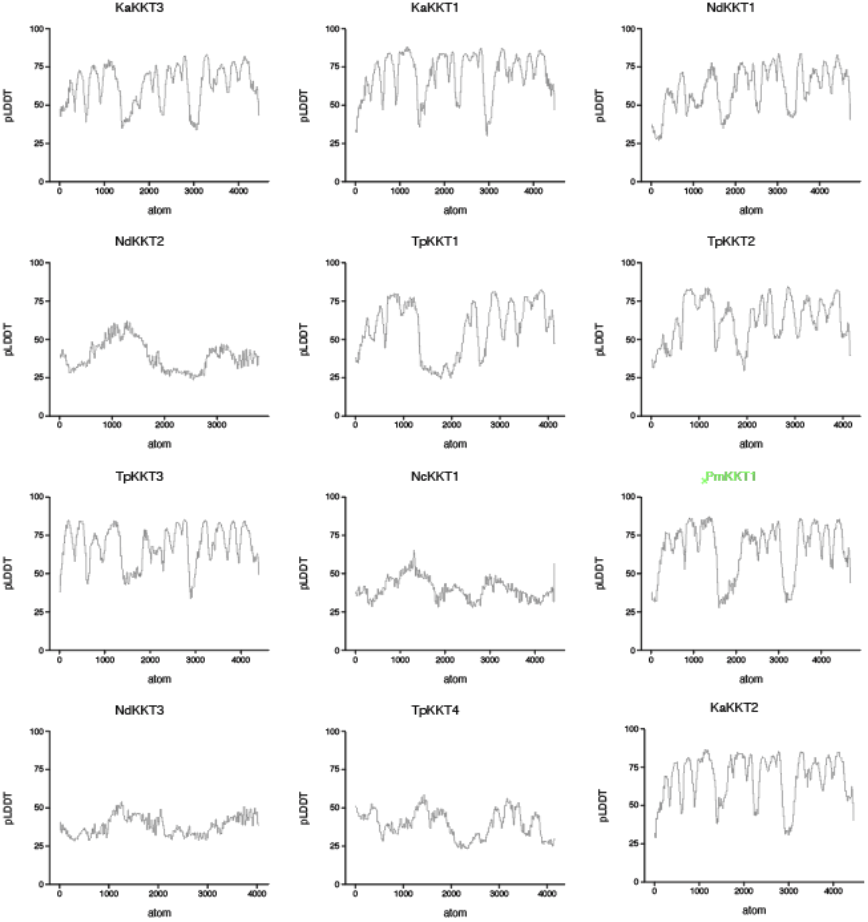
AlphaFold confidence scores for predicted killer-toxin tertiary structures. Predicted local distance difference test (pLDDT) scores for the top-ranked AlphaFold2 models of K1-like killer toxins.

**Figure S3.**
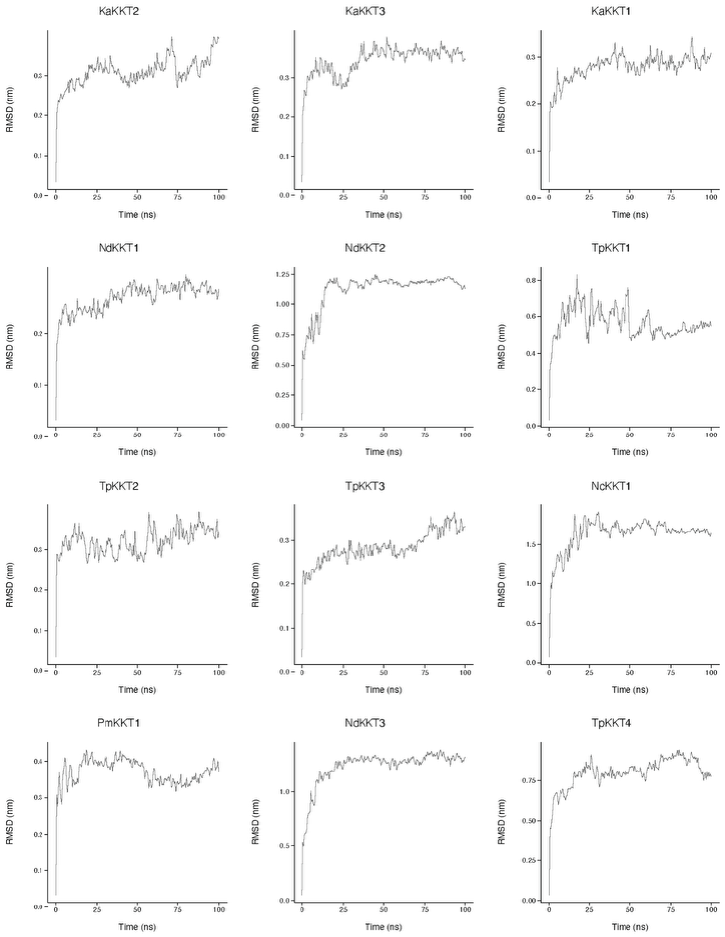
RMSD trajectories of predicted K1-like killer-toxin tertiary structures. Protein RMSD was monitored throughout 100 ns molecular dynamics simulations. Using GROMACS, each trajectory snapshot was aligned to the corresponding starting structure at 0 ns.

**Figure S4.**
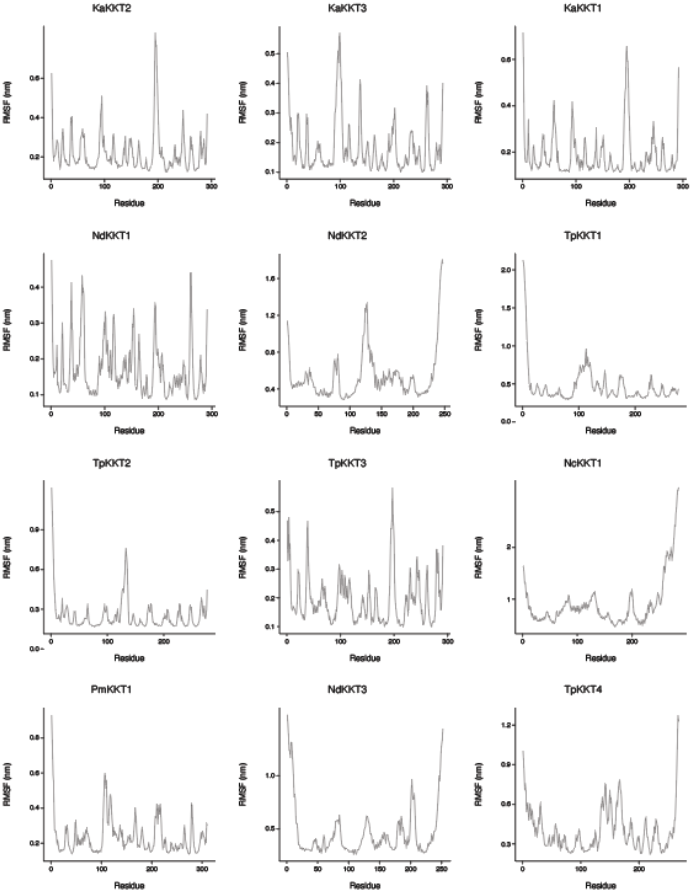
RMSF plots for every residue of each K1-like killer toxin. Protein RMSF was monitored throughout 100 ns molecular dynamics simulations for each residue of each K1-like killer toxin.

Overall, the tertiary structure models of the K1 homologs had a similar organization, which was largely consistent with their similar secondary structures. The alpha domain consists predominantly of two beta sheets followed by a two-helical bundle that is wrapped by a discontinuous, antiparallel beta sheet assembled from alpha, gamma, and beta domains. Disulfide bonds in K1 that were predicted by prior empirical studies were also predicted in the tertiary structures of K1 homologs, including one intradomain bond in the alpha (12/13) and beta (8/13) domains, as well as one interdomain bond connecting the alpha and beta domains (9/13), Along with similarities to K1, there were also differences in the number of cysteines and additional predicted disulfide bonds (Figure 3). The structures of NdKKT2, NdKKT3, and NcKKT1 were unusual in that the gamma and beta domains were mostly alpha-helical, which is different from the predominance of beta sheets observed in K1 and other homologs. Despite the poor confidence in these models, the predicted alpha domains of NdKKT2, NdKKT3, and NcKKT1 had hydrophobic helices similar to K1 and other homologs (Figure S1 and Table S5).

Alignment of the tertiary structure models confirmed that 50% of Kex cleavage sites were on flexible loops exposed to the aqueous environment, which increase to 71% after removing low-confidence models (NcKKT1, NdKKT2, and NdKKT3). Importantly, across all tertiary structure models, 75% of Kex cleavage sites aligned with the positioning of known K1 sites, including predicted cleavage sites that were not identified by the alignment of primary protein sequences. Overall, we find that the majority of K1 homologs have a similar pattern of Kex cleavage sites to K1. These data support the assignment of alpha, beta, gamma, and delta domains for 12 K1 homologs.

### Alpha domains from K1 homologs are toxic when expressed by Saccharomyces cerevisiae

Four K1 homologs are active antifungal killer toxins when expressed by *Saccharomyces* yeasts (Figure 2). The lack of antifungal activity in other K1-like killer toxins with apparently intact alpha, gamma, and beta domains could be due to a variety of factors related to ectopic expression, intrinsic killer toxin resistance of yeasts assayed for toxin sensitivity, or loss of killer toxin function due to mutation. As recombinant expression of the K1 alpha domain in *S. cerevisiae* is highly toxic, this approach was used to test the toxicity of predicted alpha domains from K1-homologs. Moreover, 12/13 K1 homologs contained at least one transmembrane region in the N-terminal half of the protein, with ten predicted to form pores, consistent with the proposed ionophoric mechanism of K1 (Figure 4 and Table S4). Cloning of alpha domains in the bacteria *Escherichia coli* was generally unsuccessful due to their surprising toxicity. Therefore, alpha domain expression plasmids were constructed using yeast recombineering by homologous recombination using *S. cerevisiae*. To test the cytotoxic function of diverse alpha domains, six were successfully cloned from K1L, KaKKT2, KaKKT3, PmKKT1, NdKKT1, and NdKKT2 and expressed in *S. cerevisiae* under the control of a galactose-inducible promoter. Five clones of each alpha domain were grown on dextrose and did not cause significant toxicity (Figure 4). The induction of the K1 alpha domain expression by galactose was toxic and prevented *S. cerevisiae* growth, whereas the induction of GFP was non-toxic (Figure 4). All other alpha domains tested were also acutely toxic to *S. cerevisiae* with little to no growth observed on galactose-containing plates.

**Figure 4 -.**
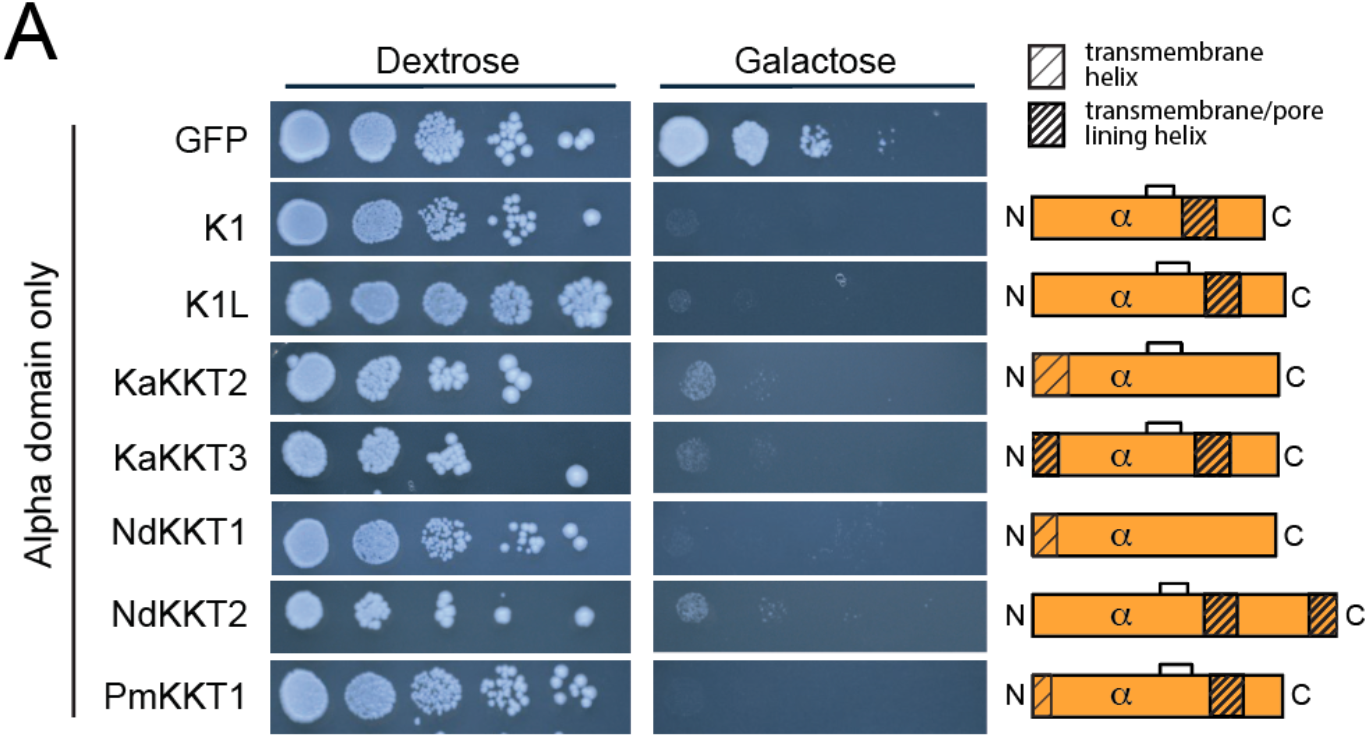
Alpha domains from K1 homologs are toxic when expressed by *S. cerevisiae*. Spot dilution assays of *S. cerevisiae* transformed with plasmids encoding the alpha domains of K1 homologs and K1 on media containing either galactose or dextrose. GFP was used as a negative control for normal yeast growth. Orange bars represent the relative length of each alpha domain and the positioning of predicted intramolecular disulfide bonds and transmembrane and pore-lining helices. All images are representative (n=5).

These data agree well with prior experiments showing that active killer toxins had cytotoxic alpha domains when expressed by *S. cerevisiae* (K1L, KaKKT2, NdKKT1, PmKKT1). Expression of the alpha domains of NdKKT2 and KaKKT3 was also toxic to *S. cerevisiae*, which was surprising given that neither was previously found to have antifungal activity. Despite the unusual abundance of helices in the gamma and beta domains of NdKKT2, the alpha domain appears to be toxic, suggesting that this protein is potentially an active killer toxin.

### Kex cleavage sites of K1-like toxins are required for antifungal activity and posttranslational cleavage

KaKKT2 was selected for analysis of toxin processing by Kex proteases and the protein was modified with a C-terminal HA epitope tag. Galactose-inducible expression enabled the detection of KaKKT2 by western blotting in whole cell lysates of *S. cerevisiae* and the presence of the uncleaved preprotoxin at ∼36 kDa in non-reducing conditions (Figure 5). Analysis of the extracellular media in non-reducing conditions showed that KaKKT2 has an apparent molecular weight of ∼24 kDa, which is consistent with an alpha-beta heterodimer. These observations support the processing of this killer toxin by posttranslational cleavage as has been observed for K1. Although the addition of the HA-tag resulted in a loss of antifungal activity, the tagged protein was correctly processed into the mature alpha-beta toxin and exported from the cell and can therefore be used to investigate the validate the predicted locations of Kex cleavage sites.

**Figure 5.**
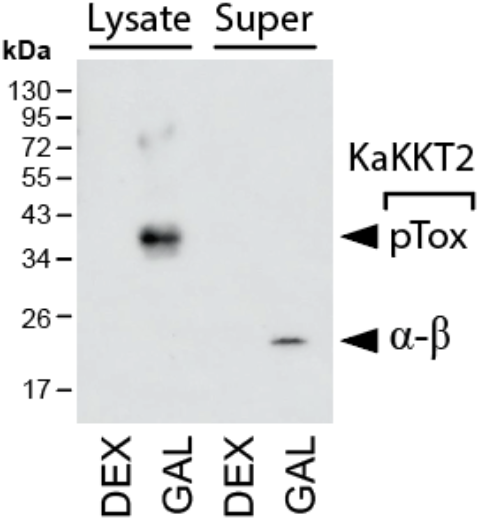
Extracellular K1-like killer toxins are processed heterodimers derived from an intracellular preprotoxin. Western blot detection of KaKKT2-HA expressed by *S. cerevisiae* in both extracellular medium (super) and whole-cell lysate (lysate), with detection of the preprotoxin (ppTox) and the mature alpha-beta heterodimer (α-β) in non-reducing conditions. The beta domain (β) is observed under reducing conditions. All images are representative (n>3).

### Kex cleavage sites of K1-like toxins are required for antifungal activity and posttranslational cleavage

Mutations at the Kex cleavage sites of K1 prevent toxin maturation, resulting in inactivation of the exogenously expressed toxin. If cleavage occurs at the predicted kex cleavage sites in the KKTs and is required for the maturation of the toxin, we would expect that mutations at these sites would produce inactive toxins. To abolish the posttranslational cleavage of KaKKT2, PmKKT1, NdKKT1 by Kex proteases, alanine mutations were made at each of the putative cleavage sites and were tested for activity with killer assays. Toxin expression was again controlled by a galactose-inducible promoter and, in contrast to the large halos of growth inhibition of the wild-type toxins, Kex site mutants in all three toxins showed complete loss of toxin activity (Figure 6A).

**Figure 6.**
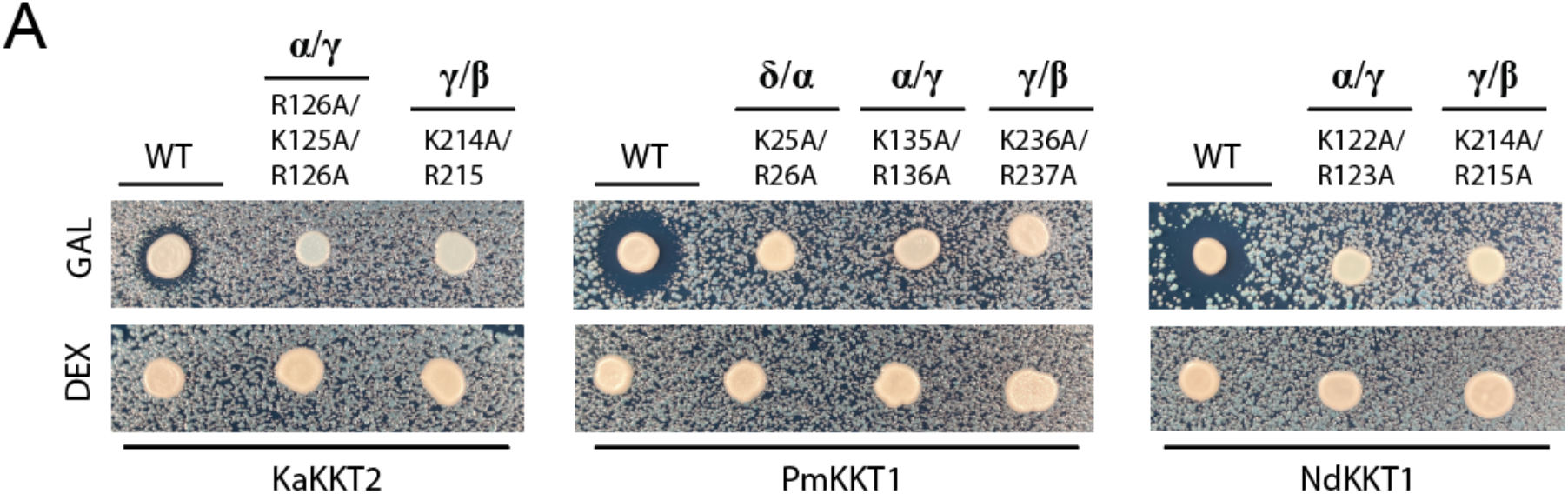
Loss of Kex cleavage sites in K1 homologs leads to inactivation and failure of posttranslational cleavage of protoxin. (A) In three K1-like killer toxins basic residues at putative Kex cleavage sites were mutated to alanine and then tested for antifungal activity on agar compared to the wild-type protein.

Together, these results support the identification of protein products corresponding to the major processing events of KaKKT2, including signal peptidase cleavage and Kex cleavage sites predicted to delineate functional toxin domains. The ability to predict Kex cleavage sites based on amino acid sequence and protein structure in KaKKT2 support the predictions of processing sites in other K1-family killer toxins.

## Discussion

This study demonstrates that K1 homologs likely require the carboxypeptidases Kex1 and Kex2 for posttranslational maturation and activation of their antifungal activities. These results parallel the well-known requirement of Kex enzymes for the maturation of the canonical K1 toxin from *Saccharomyces cerevisiae* and reinforce the evolutionary conservation of Kex-dependent processing pathways in killer toxins across different yeast species. The conservation of Kex cleavage sites in K1 homologs supports their broader role in killer toxin maturation. Multiple sequence alignment revealed that conserved basic sequences define domain boundaries between the delta, alpha, gamma, and beta domains of K1 homologs. These conserved sequences—often characterized by pairs of lysine and arginine residues—likely function as recognition sites for Kex1 and Kex2, facilitating the cleavage and maturation of killer toxins. This is particularly evident in the predicted cleavage sites at the delta-alpha, alpha-gamma, and gamma-beta boundaries, where conserved basic residues were identified across multiple K1 homologs.

Interestingly, despite the conservation of these cleavage motifs, some homologs showed variation in the position of the cleavage sites relative to K1. For example, while the Kexa^/g^ site was conserved in most homologs, its position was shifted toward the N-terminal domain compared to K1. This variation suggests that while the general requirement for Kex-dependent cleavage is conserved, the exact cleavage positions may have evolved to accommodate differences in domain architecture or functional requirements among K1 homologs. Tertiary structure predictions generated using AlphaFold2 revealed a similar organization of alpha, beta, gamma, and delta domains across all homologs. These models found that most predicted Kex cleavage sites are located in flexible, exposed regions of the protein, which generally overlapped with the known K1-a cleavage sites in tertiary structure models, even when sites did not overlap using multiple sequence alignments. The positioning of these sites on loops situated between elements of secondary structure provides accessibility for Kex1 and Kex2 to cleave the protein during maturation.

The cytotoxicity assays further demonstrated the importance of the alpha domain in determining the antifungal activity of K1 homologs. When expressed in *S. cerevisia*e, several K1 homolog alpha domains (K1L, KaKKT2, NdKKT1, and PmKKT1) exhibited strong cytotoxicity, consistent with their previously identified killer toxin activity. Interestingly, the alpha domains of NdKKT2 and KaKKT3, which did not display antifungal activity in their full-length forms, were also cytotoxic when expressed independently. This suggests that factors other than alpha domain toxicity, such as proper folding, domain interaction, or target specificity, may be required for the full-length toxins to exert their antifungal effects.

These findings open several avenues for further investigation. One possibility is that differences in the processing, folding, or stability of full-length killer toxins could explain the lack of activity in certain homologs, despite their toxic alpha domains. Additionally, the identification of homologs with cytotoxic alpha domains that lack detectable killer activity in their full-length forms raises questions about the evolution and functional divergence of killer toxins. Future work should explore the role of other domains in modulating toxin activity and investigate the environmental or evolutionary pressures that drive the conservation of Kex cleavage sites and toxin domain architecture.

In conclusion, this study highlights the critical role of Kex-dependent processing in the maturation and activation of K1 homologs, demonstrating both structural and functional conservation across diverse yeast species. The identification of conserved dibasic cleavage sites and the functional characterization of alpha domains in K1 homologs provide valuable insights into the molecular mechanisms underlying killer toxin activity. These findings lay the groundwork for future studies aimed at elucidating the broader functional diversity of killer toxins and their potential applications in antifungal therapies or industrial biotechnology.

## Methods

### Yeast transformation

Plasmids were transformed into *S. cerevisiae* using the lithium acetate method. Transformants containing pAG426-derived constructs were selected on complete minimal medium lacking uracil (CM -ura) and containing dextrose. Plates were incubated at 30°C for 48 h.

### KKT cloning and mutagenesis

K1-alpha constructs were built by homologous recombination in *S. cerevisiae*. DNA fragments encoding killer toxins and the plasmid backbone (pAG426-GAL-ccdB) were created by PCR using primers with 30 bp of overlapping homology. *S. cerevisiae* was transformed with the PCR products using the LiAc method, and transformants were selected using CM -ura and grown at 25°C. KKT genes were cloned as described previously.

### Induction of plasmid-encoded killer toxins

Plasmid-containing clones were inoculated into 2 mL liquid CM -ura medium containing dextrose and cultured overnight at 30°C with shaking at 200 rpm. Following overnight growth, 1 mL of culture was transferred directly into 24 mL CM -ura medium containing dextrose or washed three times with 0.5 mL phosphate-buffered saline (PBS; pH 7.0), suspended in CM - ura medium containing galactose, and transferred into 24 mL CM -ura medium containing galactose. Cultures were grown overnight at 30°C with shaking at 200 rpm. To collect culture supernatant, cultures were centrifuged at 2,432 x g for 5 min. Cell pellets were suspended in 1 mL fresh medium. The culture supernatants were retained for subsequent analysis of extracellular toxin.

### Yeast cell lysis

Cell pellets (∼2 × 10^8^ viable cells) were suspended in an equal volume of lysis buffer (50 mM Tris, 100 mM NaCl, and 1 mM EDTA (pH 7.5)). EDTA-free complete protease inhibitor (Roche), prepared in 10 mL deionized water, was added at a 1:10 ratio relative to the lysis buffer. Glass beads were added at a volume equal to that of the lysis buffer, producing a 1:1:1 ratio of cells, lysis buffer, and glass beads. Cell suspensions were vortexed for 2 min at 4°C and rested on ice for 2 min. This cycle was repeated four times. The lysates were centrifuged at 1,000 × g for 8 min at 4°C to pellet the glass beads. The supernatants were transferred to fresh 1.5 mL microcentrifuge tubes and centrifuged at 10,000 × g for 30 min at 4°C to remove residual cellular debris.

### Protein quantification

Protein concentrations were estimated from absorbance at 280 nm using a NanoDrop spectrophotometer. The instrument was blanked using lysis buffer containing protease inhibitor at the same 1:10 ratio used for sample preparation. The volume of 200 μg of total estimated protein was adjusted to 40 μL with lysis buffer. Samples were supplemented with 10 μL 5× sample buffer (300 mM Tris-HCl (pH 6.8), 10% SDS, 50% glycerol, 0.01% bromophenol blue) and heated at 100°C for 5 min.

### Culture-supernatant concentration

Culture supernatant removed from cultures with induced killer toxin was transferred to a 15 mL Amicon Ultra centrifugal filter with a 10 kDa molecular-weight cutoff and centrifuged at 2,432 × g at 4°C until the supernatant was concentrated to ∼150 μL (∼130-fold concentration). Protein samples were further concentrated by trichloroacetic acid (TCA) precipitation. Tween-20 was added to a final concentration of 0.5% (v/v), followed by 100% TCA to a final concentration of 20% (v/v). Samples were mixed thoroughly, incubated on ice for 30 min, and centrifuged at 15,000 × g for 15 min at 4°C. The supernatant was removed, and pellets washed with 300 μL ice-cold acetone. After the removal of the acetone, precipitated protein were suspended in ∼30 μL lysis buffer with protease inhibitor. Samples were then heated at 100°C for 5 min before analysis by western blotting.

### Western Blotting

For each sample, 80 μg total protein was loaded onto a 15% SDS-polyacrylamide gel and separated in Tris–glycine running buffer. Electrophoresis was performed at 80 V for 15 min. After samples entered the gel, electrophoresis was continued at 100 V for 75 min. Following electrophoresis, proteins were transferred to a nitrocellulose membrane using a TurboBlot semidry transfer system (BioRad). The transfer stack consisted of three sheets of Whatman chromatography paper, the nitrocellulose membrane, the gel, and three additional sheets of chromatography paper. All components were pre-equilibrated in transfer buffer (25 mM Tris-HCl, 192 mM glycine, and 20% (v/v) methanol at pH 8.3). Transfer was performed for 60 min at a constant voltage of 25 V, with the current limit set to 1.0 A. The measured current during transfer was approximately 0.3 - 0.5 A. Immediately after transfer, membranes were incubated in 0.1% (w/v) Ponceau S solution for 1 min at room temperature to assess protein transfer. Excess stain was removed by washing with Tris-buffered saline containing 0.1% Tween 20 (TBST). Membranes were blocked for 1 h at room temperature in TBST and 3% nonfat dry milk, with rocking at 25 rpm. Blocked membranes were incubated for 1 h at 20°C with an HRP-conjugated anti-HA antibody diluted 1:2,000 to 1:6,000 in TBST containing 0.1% Tween 20, with rocking at 25 rpm. Membranes were washed in TBST containing 0.1% Tween 20 for successive intervals of 1, 1, 2, and 4 min at 20°C with rocking at 25 rpm. The chemiluminescent signal was developed by incubating the membranes for 1 min with the two components of Cytiva ECL Select reagent mixed at a 1:1 ratio. Membranes were imaged using an ImageQuest system.

### Killer Assays

Killer assays were performed on pH 4.0 methylene blue YPD and YPG agar plates. Killers were grown overnight in 25°C CM-ura dextrose-containing liquid media, shaking at 200 rpm. *K. africana* (NCYC2729) was used as a sensitive lawn and was grown under the same conditions in YPD liquid media. The lawn was diluted to an OD600 of 0.0015, and 1 mL was spread on the plate. Killer yeast cells were centrifuged at 2,000 rpm for 1 min, and media was removed. 2 µL of killer cell slurry was spotted on top of the already seeded *K. afticana*. Plates were grown for 5-7 days at 22°C before imaging.

### Molecular Modeling methods

To generate predictions of killer toxin tertiary structures, translated protein sequences were analyzed by AlphaFold2. SignalP and TargetP were used to predict signal sequences. The AF2 models underwent 100 ns molecular dynamics simulations with GROMACs. Simulation conditions were performed as previously described. Clustered models were generated with varying RMSD cutoffs. Cutoff was determined by the number of frames in top clustered model. Cutoffs with the top model containing 55% or more frames were chosen. RMSD, and RMSF over the simulation were plotted. Ramachandran plots of the energy-minimized structure and post-simulation clustered model were made. Secondary structure predictions of homologs were also performed with Jpred.

The predicted structures were aligned in PyMOL using the super command. Using common structural motifs shared between the KKTs and K1, functional domains were identified. Cleavage by Kex2 was assumed to be shared by the KKTs, so cleavage was predicted to be at basic, preferably dibasic, sites on exposed loops. Where this ideal criterion was not met, the domain boundary was predicted at the closest basic residue to the domain boundary established by structural homology. In the case of more than two basic residues at a site, cleavage was predicted to be at the C-terminal end of the site.

### Other computational methods

Signal sequence predictions were made using SignalP (v6.0) and targetP (v2.0). ChatGPT Codex was used to confirm the accuracy of manually curated datasets kex cleavage site datasets from fasta formatted protein sequences of all K1 homologs.

### Assay for alpha domain toxicity

Yeast expressing the alpha domains of different killer toxins were grown overnight with shaking at 200 rpm in CM-ura dextrose media at 25°C. *S. cerevisiae* K1-alpha domain and GFP were included as controls. Overnight cultures were diluted to an OD600 of 1.5 and serially diluted tenfold to 1 × 10^−5^. Dilutions were inoculated on plates with dextrose or galactose with a metal pinning tool. Plates were grown at 25 °C for 5-7 days before imaging.

---

**Table S1** SignalP and TargetP signal peptidase cleavage site predictions

**Table S2** pLDDT scores for AlphaFold2 predictions of K1 homologs

**Table S3** Ramachandran values of tertiary structure models pre- and post-MD simulation.

**Table S4** Location of predicted pore-forming and membrane-interacting helices

**Table S5** Plasmids used in this study. pEK006 (pAG426-Gal-K1)

pML117 (pAG426-Gal-KaKKT2)

pML116 (pAG426-Gal-PmKKT1)

pML115 (pAG426-Gal-NdKKT1) pAG426-Gal-GFP

pSC048 (pAG426-Gal-KaKKT2-HA)

pSC049 (pAG426-Gal-KaKKT2 SDM1-HA)

pSC050 (pAG426-Gal-KaKKT2 SDM1+2-HA)

**Table S6** Primers used in this study.

File S1. Protein sequences of all killer toxins used in this study.

File S2. DNA sequence of all plasmids used in thai study.

