## Supplementary Tables for "Defining the domain organization of K1-like killer toxins"

| | | # SignalP-6.0 | | | TargetP 2.0 | | |
|---|---|---|---|---|---|---|---|
| # ID | Prediction | OTHER | SP(Sec/SPI) | CS Position | OTHER | SP(Sec/SPI) | CS Position |
| K1 | SP | 0.24093 | 0.759073 | CS pos: 36-37. Pr: 0.5257 | 0.0536 | 0.9463 | **CS pos: 26-27. VVA-LN. Pr: 0.8208** |
| K1L | SP | 0.000152 | 0.999866 | CS pos: 26-27. Pr: 0.9815 | 0.00257 | 0.997316 | CS pos: 26-27. TSG-AD. Pr: 0.8724 |
| KaKKT2_118k.af | SP | 0.000154 | 0.999842 | CS pos: 18-19. Pr: 0.9825 | 0.002186 | 0.997813 | CS pos: 18-19. ALA-AF. Pr: 0.8178 |
| KakKT3_117k.af | SP | 0.000174 | 0.999845 | CS pos: 18-19. Pr: 0.9803 | 0.002218 | 0.99778 | CS pos: 18-19. ALA-AF. Pr: 0.8174 |
| KaKKT1 | SP | 0.000156 | 0.999843 | CS pos: 18-19. Pr: 0.9805 | 0.004674 | 0.995322 | CS pos: 18-19. ALA-AF. Pr: 0.8221 |
| NdKKT1_XP_003 | NO_SP | 0.780801 | 0.219191 | - | 0.126445 | 0.87338 | **CS pos: 16-17. VQA-AF. Pr: 0.7298** |
| NdKKT2 | SP | 0.000224 | 0.999782 | CS pos: 24-25. Pr: 0.9310 | 0.000868 | 0.999044 | **CS pos: 19-20. SLA-GM. Pr: 0.4350** |
| TpKKT1_XP_003 | SP | 0.000155 | 0.999842 | CS pos: 14-15. Pr: 0.9802 | 0.101536 | 0.898463 | CS pos: 14-15. TVA-DD. Pr: 0.9357 |
| TpKKT2_XP_003 | SP | 0.000159 | 0.999838 | CS pos: 14-15. Pr: 0.9792 | 0.117272 | 0.882726 | CS pos: 14-15. TIA-DD. Pr: 0.9526 |
| TpKKT3_XP_003 | SP | 0.250237 | 0.749777 | CS pos: 19-20. Pr: 0.7810 | 0.000222 | 0.999777 | CS pos: 19-20. ADA-SF. Pr: 0.7520 |
| NcKKT1_XP_003 | SP | 0.000165 | 0.999833 | CS pos: 18-19. Pr: 0.9841 | 0.000455 | 0.999532 | CS pos: 18-19. TLA-SD. Pr: 0.8437 |
| PmKKT1 | SP | 0.324883 | 0.675139 | CS pos: 19-20. Pr: 0.6965 | 0.005387 | 0.994559 | CS pos: 19-20. VAA-TP. Pr: 0.8092 |
| NdKKT3 | SP | 0.168753 | 0.831231 | CS pos: 19-20. Pr: 0.8150 | 0.013604 | 0.986322 | CS pos: 19-20. SLA-DM. Pr: 0.7325 |
| TpKKT4_XP_003 | NO_SP | 1.000005 | 0 | - | 0.892759 | 0.107147 | - |

Table S1. Signal peptidase cleavage prediction by Signal P and Target P.

| Name | Signal peptide | Delta | Alpha | Gamma | Beta | Total |
|---|---|---|---|---|---|---|
| K1 | 36 | 8 | 105 | 84 | 83 | 316 |
| K1L | 26 | 10 | 111 | 101 | 92 | 340 |
| KaKKT2 | 18 | - | 108 | 91 | 93 | 310 |
| KaKKT3 | 18 | - | 108 | 91 | 93 | 310 |
| KaKKT1 | 18 | - | 109 | 90 | 93 | 310 |
| NdKKT1 | - | 25 | 98 | 92 | 92 | 307 |
| NdKKT2 | 24 | 16 | 134 | 76 | 21 | 271 |
| TpKKT1 | 14 | - | 99 | 80 | 98 | 291 |
| TpKKT2 | 14 | - | 99 | 80 | 98 | 291 |
| TpKKT3 | 19 | - | 109 | 90 | 92 | 310 |
| NcKKT1 | 18 | 21 | 129 | 94 | 41 | 303 |
| PmKKT1 | 19 | 7 | 110 | 101 | 92 | 329 |
| NdKKT3 | 19 | 21 | - | - | - | 271 |
| TpKKT4 | - | - | 143 | 88 | 37 | 268 |

Table S2. Domain boundaries of K1-like killer toxins.

| | pLDDT AF2 | pLDDT AF3 | difference | percent |
|---|---|---|---|---|
| K1 | 51.5318 | 31.92532 | 19.60648 | 0.614136 |
| K1L | 54.88956 | 48.31668 | 6.57288 | 0.136037 |
| KaKKT1 | 72.42544 | 53.6901 | 18.73534 | 0.348953 |
| KaKKT2 | 69.89803 | 68.75305 | 1.14498 | 0.016654 |
| KaKKT3 | 64.39909 | 55.21927 | 9.17982 | 0.166243 |
| NcKKT1 | 39.30685 | 40.92341 | -1.61656 | -0.0395 |
| NdKKT1 | 62.08369 | 51.6114 | 10.47229 | 0.202907 |
| NdKKT2 | 38.27589 | 54.90255 | -16.6267 | -0.30284 |
| NdKKT3 | 37.18452 | 53.17854 | -15.994 | -0.30076 |
| PmKKT1 | 67.66196 | 41.68782 | 25.97414 | 0.623063 |
| TpKKT1 | 58.18974 | 50.81374 | 7.376 | 0.145158 |
| TpKKT2 | 64.18728 | 40.6278 | 23.55948 | 0.579886 |
| TpKKT3 | 70.18415 | 46.99225 | 23.1919 | 0.493526 |
| TpKKT4 | 38.44608 | 47.66059 | -9.21451 | -0.19334 |
| MIN | 37.18452 | 31.92532 | 7.31154 | 0.177866 |
| MAX | 72.42544 | 68.75305 | 25.97414 | 0.623063 |
| average | 56.33314857 | 49.02160857 | 7.311538571 | 0.1778659286 |

Table S3 – pLDDT scores from AlphaFold predictions of K1-like killer toxins

| | pre-MD rama | post-MD rama |
|---|---|---|
| KaKKT1 | 92.76 | 91.57 |
| KaKKT2 | 91.72 | 91.95 |
| KaKKT3 | 88.62 | 89.22 |
| NcKKT1 | 87.28 | 90.08 |
| NdKKT1 | 88.2 | 90.98 |
| NdKKT2 | 92.65 | 86.92 |
| NdKKT3 | 92 | 86.83 |
| PmKKT1 | 90.26 | 91.88 |
| TpKKT1 | 85.09 | 92.8 |
| TpKKT2 | 93.09 | 89.02 |
| TpKKT3 | 92.04 | 89.53 |
| TpKKT4 | 83.83 | 88.31 |
| Average | 89.795 | 89.92417 |

Table S4 – Ramachandran favorable areas pre- and post-MD simulation.

| Name | Transmembrane helices | Pore lining helicies |
|---|---:|---:|
| K1 | 111-126 | 111-126 |
| K1L | 112-127 | 112-127 |
| KaKKT2 | 18-33 | - |
| KaKKT3 | 14-34,90-105 | 14-34,90-105 |
| KaKKT1 | 18-33,90-105 | 90-105 |
| NdKKT1 | 14-32 | - |
| NdKKT2 | 116-131,162-179,186-201,235-250 | 116-131,162-179,235-250 |
| TpKKT1 | 43-58 | - |
| TpKKT2 | 44-59 | - |
| TpKKT3 | 96-111 | 96-111 |
| NcKKT1 | 113-128,155-170,181-196,229-246 | 113-128,155-170,229-246 |
| PmKKT1 | 18-33,103-118 | 103-118 |
| NdKKT3 | 41-56,118-133,162-179,186-201,235-250 | 118-133,162-179,235-250 |
| TpKKT4 | 23-38,108-123 | 23-38,108-123 |

Table S5 – Position of hydrophobic helices in K1-like killer toxins.

| Primer | Sequence | Description |
|---|---|---|
| prSC001 | cgtcgacctcgagtcatgta | ccdB Gal pAG426 recombineering backbone 2 fwd |
| MS048 | cagaatgggcagacattacg | primer set 2: ura3 portion |
| MS049 | gttacttcttctgccgcctg | primer set 3: ura3 going 5' to 3' dir. |
| MS065 | ccggggttttttctccttgacgttaaagtatagag | primer set 2: before gene of interest |
| prSC002 | ctctatactttaacgtcaaggagaaaaaaccccggatg TGGAGTACTTCGAACACCTATCATGC | Recombineering K1L alpha fwd pSC001 pAG |
| prSC003 | gtgacataactaattacatgactcgaggtcgacgtca CCTCTTCAGAGGTTGATCACTAGTG | Recombineering K1L alpha rev pSC001 pAG |
| prSC010 | ctctatactttaacgtcaaggagaaaaaaccccggatg GAACCTAATCCATCTGTTTTGACTCTAG | Recombineering NdKKT2 (pML113) alpha forward |
| prSC011 | gtgacataactaattacatgactcgaggtcgacgtca TTTTACCCAAACTGGCAAAGGTCC | Recombineering NdKKT2 (pML113) alpha reverse |
| prSC014 | ctctatactttaacgtcaaggagaaaaaaccccggatg ATATTTCCATTTTACGACTTTATACACAC | Recombineering PmKKT1 (pML116) alpha forward |
| prSC015 | gtgacataactaattacatgactcgaggtcgacgtca AACAGGAGAACCATTTCCGTCG | Recombineering PmKKT1 (pML116) alpha reverse |
| SC019 | TGGGGTAATTAATCAGCGAAGC | alpha amplification universal pAG426-gal FORWARD |
| SC020 | GACTTCAGGTTGTCTAACTCCTTCC | alpha amplification universal pAG426-gal REVERSE |
| SC021 | AATACTTTCAACATTTTCGGTTTG | sequencing alpha universal pAG426-gal |
| SC040 | ctctatactttaacgtcaaggagaaaaaaccccggatg GAAGCGCCGTGGTATGACAAG | K1-a recombineering forward |
| SC041 | gtgacataactaattacatgactcgaggtcgacgtca AGCAACGATAGCGCCATTAGG | K1-a recombineering reverse |
| SC042 | ctctatactttaacgtcaaggagaaaaaaccccggatg gcatttccatattatgattatattcatttagg | KaKKT2/KaKKT3 alpha recombineering fwd |
| SC043 | gtgacataactaattacatgactcgaggtcgacgtca tctctttcggagagctgttgg | KaKKT2 alpha recombineering rev |
| SC044 | gtgacataactaattacatgactcgaggtcgacgtca tctctttcggagagccgttgg | KaKKT3 alpha recombineering rev |

Table S6 – Primers used in this study
